# Transcriptional antitermination integrates the expression of loci of diverse phage origin in the chimeric *Bartonella* Gene Transfer Agent BaGTA

**DOI:** 10.1101/2024.11.20.624540

**Authors:** Aleksandr Korotaev, Quirin Niggli, Valeria Congedi, Christoph Dehio

## Abstract

Gene Transfer Agents (GTAs) are mobile genetic elements derived from bacteriophages that mediate genome-wide horizontal gene transfer (HGT) in diverse groups of prokaryotes. BaGTA, encoded by all the pathogens of the genus *Bartonella*, is a chimeric GTA that evolved by the domestication of two phages. The run-off-replication module ROR of one phage is integrated with the capsid production, DNA packaging and lysis machinery Bgt of a second phage. Restricted to a self-sacrificing subset of the bacterial population, the position-specific DNA amplification and packaging of a genomic plasticity region enriched for genes involved in host interaction and adaptation selectively enhances the HGT frequency of these pathogenicity genes. This feature of BaGTA is considered a key innovation underlying the evolutionary success of *Bartonella*. Little is known, however, about the mechanism mediating the coordinated expression of the *ror* and *bgt* loci. Here, we established the regulatory hierarchy, with *ror* acting upstream of the capsid gene cluster *bgtA-K*. BrrG, encoded by the *ror* locus, controls the transcription of the *bgtA-K* operon by functioning as a processive antiterminator. This study provides the first insights into the mechanism controlling the coordinated expression of the two BaGTA modules of divergent phage origin. Beyond BaGTA, we propose that antitermination is a broadly relevant mechanism for controlling HGT by GTAs of the Alphaproteobacteria.

## Introduction

Timed and tightly controlled gene expression is crucial in bacterial cells. A variety of mechanisms to control transcription are known, comprising both positive and negative regulators (reviewed in Turnbough, 2019). One of the mechanisms of negative regulation involves transcription attenuation at termination sites (reviewed in Santangelo and Artsimovitch, 2011). The regulation of transcription attenuation is achieved by a diverse class of antiterminators which counteract the attenuation (Santangelo and Artsimovitch, 2011). Antiterminators fall into two categories and can either act specifically at a single, defined terminator site or convert RNA polymerase into a termination-resistant complex that is generally resistant to multiple termination sites over a longer distance (Santangelo and Artsimovitch, 2011; reviewed in Goodson and Winkler, 2018). In the latter case, such antiterminators are referred to as processive antiterminators. Well-known examples of processive antiterminators are Q and N of bacteriophage lambda which control progression of the infection cycle (Goodson and Winkler, 2018).

Peculiar examples of phage evolution are Gene Transfer Agents (GTAs) (reviewed in Québatte and Dehio, 2019; Banks and Tung, 2024; Craske et al., 2024). GTAs are mobile genetic elements, which emerged from defective temperate bacteriophages. GTAs encode bacteriophage-like particles and in contrast to phages transfer unspecific pieces of host bacterial DNA. Despite their phage origin, GTAs are not self-transmitting because the DNA packaging capacity of a GTA particle is too small to package the entire gene cluster encoding it. Therefore, GTAs are transferred vertically (Banks and Tung, 2024; Craske et al., 2024; Kogay et al., 2022; Lang et al., 2012). GTAs are found both in bacteria and archaea, and are one of the non-canonical mechanisms of horizontal gene transfer (HGT) (Arnold et al., 2022).

Bacterial pathogens of the genus *Bartonella* encode a chimeric GTA known as BaGTA (Québatte and Dehio, 2019; Berglund et al., 2009; Guy et al., 2013). BaGTA encodes the gene clusters *bgtA-K,* responsible for the synthesis and assembly of GTA particles, *bgtL-R*, with orphan functions, and *bgtS-V*, involved in putative host cell lysis (Fig. 1A). DNA packaging in GTA particles is coupled to a process of site-specific DNA amplification mediated by the *ror* locus located in between the *bgtL-R* and *bgtS-V* gene clusters. This locus, emerging from a different phage domestication event, includes an origin of run-off replication (ROR) and the associated *brr* gene cluster *(brrA-G)* (Berglund et al., 2009; Guy et al., 2013; Québatte et al., 2017; Tamarit et al., 2018) (Fig. 1A). The *bgtA-K* and *ror* loci are located within a high plasticity zone of the genome that is prone to DNA integrations and is highly enriched for genes encoding virulence properties involved in host interaction and adaptation (Berglund et al., 2009; Guy et al., 2013). This specific genome architecture leads to ROR-driven amplification and packaging of these pathogenicity genes, resulting in elevated transfer rates to other *Bartonella* bacteria compared to housekeeping genes encoded more distantly to the ROR origin (Québatte et al., 2017). This process is considered to provide the recombination substrates for the duplication and evolutionary diversification of genes encoding the type IV secretion systems (T4SSs) and cognate effector proteins which collectively represent a key innovation associated with the adaptive radiation of these pathogens to diverse mammalian hosts (Guy et al., 2013; Engel et al., 2011). Additionally, it is also considered to facilitate the fast evolution of surface-located autotransporters serving in host interaction and adaptation, such as the recombination-driven antigenic variation of the CFA autotransporter (Siewert et al., 2022). Thus, BaGTA is a crucial factor for the fitness and adaptability of pathogens in the *Bartonella* genus (Québatte and Dehio, 2019). However, little is known about how the expression of the *bgt* and *brr* loci derived from separate phages is regulated in a coordinated manner.

**Figure 1.**
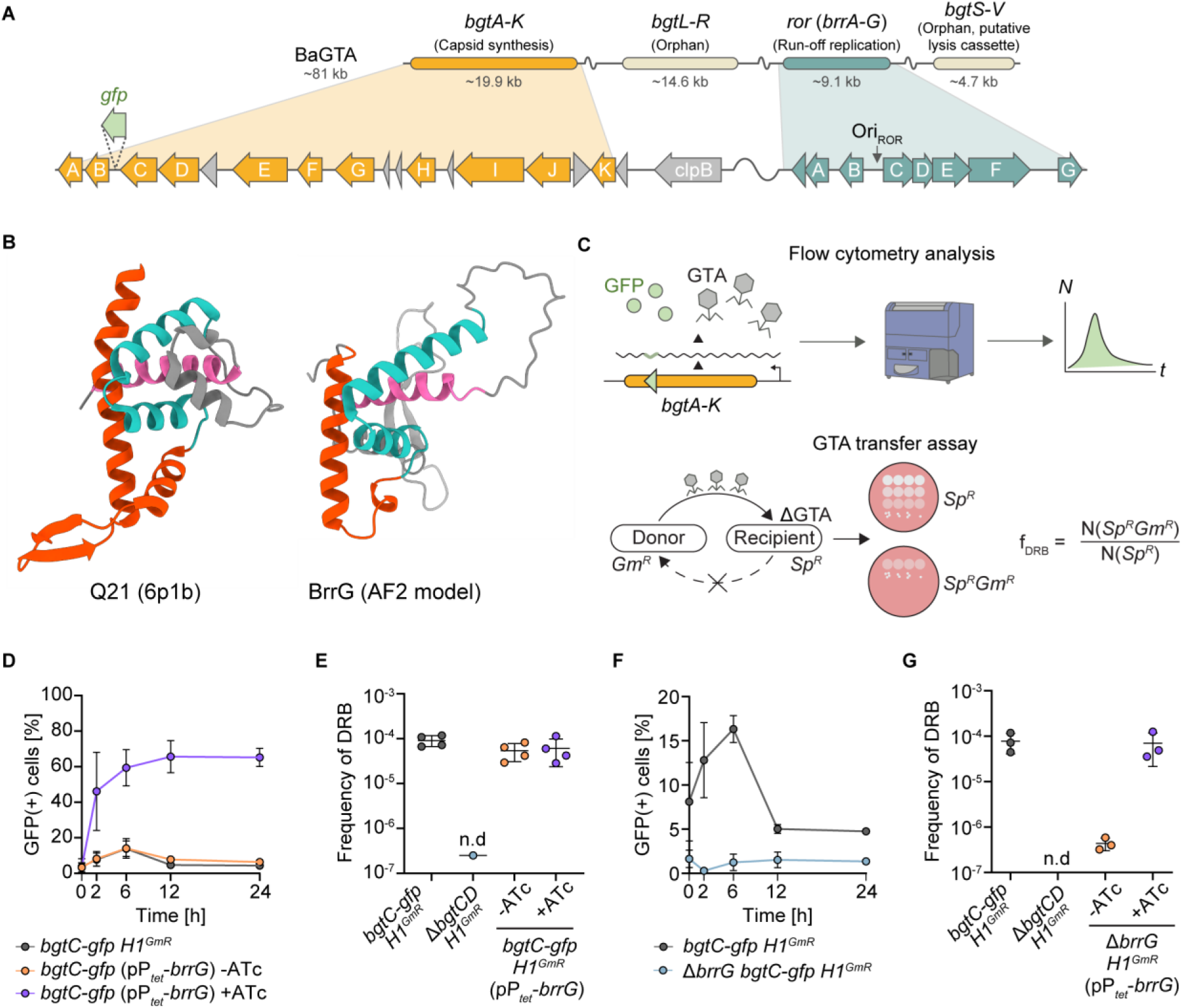
BrrG encodes a putative antiterminator essential for *bgtA-K* expression and BaGTA-mediated gene transfer. **(A)** Schematic representation of the BaGTA locus. Upper part: Structure and size of the *bgtA-K* and *ror* gene clusters. Lower part: Close-up of the *bgtA-K* locus encoding capsid and packaging functions and of the *ror* locus, encoding the run-off-replication origin *ori_ROR_* and the associated *brrA-G* gene cluster. For the reporter strain *bgtC-gfp* and its derivatives used in D-G, the insertion site of the transcriptional *gfp* reporter within the *bgtA-K* locus is indicated. **(B)** Comparison of the structure of Q21 antiterminator (6p1b) of phage P21 and Alphafold2 model of BrrG (A0A0R4J952). Similar structural features are presented in identical colour. **(C)** Schematic presentation of (i) the flow cytometry assay to monitor expression of the *bgtA-K* locus using the *bgtC-gfp* transcriptional reporter strain (upper panel) and (ii) the assay to monitor GTA transfer via transfer of *Gm^R^* from the GTA-positive donor strain to the *ΔbgtCD Sp^R^* GTA-negative (ΔGTA) recipient strain resulting in *Sp^R^Gm^R^*bacteria and formula to calculate the frequency of double resistant bacteria (f_DRB_) as a proxy for GTA transfer. **(D)** BrrG overproduction increases the bacterial population expressing *bgtA-K*. BrrG expression was induced in strain *bgtC-gfp* (pP_tet_-*brrG*) with anhydrotetracycline (aTc). Strain *bgtC-gfp H1^GmR^* (*bgtC-gfp BH01490::Himar1^GmR^)* displaying wild-type properties served as a positive control. **(E)** BrrG overproduction does not enhance GTA transfer. Strain *bgtC-gfp H1^GmR^*displaying wild-type properties and the GTA-negative mutant Δ*bgtCD H1^GmR^*(*ΔbgtCD BHRS00695::Himar1^GmR^*) served as a positive and negative control, respectively. **(F)** Deletion of *brrG* (Δ*brrG bgtC-gfp H1^GmR^* [*ΔbrrG BH01490::Himar1^GmR^*]) results in reduced expression of the *bgtA-K* locus. **(G)** Overproduction of BrrG in a Δ*brrG* mutant background restores GTA transfer frequency to the wild-type level. BrrG expression was induced in strain Δ*brrG H1^GmR^*(pP_tet_-*brrG*) with aTc. **(D-G)** The time-course experiments were started with the incubation of CBA agar-grown bacteria in M199 medium supplemented with 10% FCS. 25 ng/ml aTc was added where indicated to induce expression by the P*_tet_* promoter. The experiments were performed in biological triplicates (D, F, G) or quadruplicates (E), error bars indicate SD.

In this study, we elucidated the regulatory hierarchy between the *bgtA-K* and *ror* loci, resolving the mechanism of gene expression control. We discovered that BaGTA operates under a strict regulatory framework, with a protein encoded by the *ror* locus and controlling the expression of the *bgtA-K* capsid-production locus. Specifically, our research revealed that in absence of BaGTA activation, transcription of the *bgtA-K* locus is attenuated by a putative terminator site. Upon BaGTA activation by a yet unknown process, the *ror* locus-encoded BrrG overcomes this transcription attenuation of the *bgtA-K* locus by acting as a processive antiterminator. Furthermore, we propose that antitermination is a common mechanism of GTA regulation in the Alphaproteobacteria. Collectively, these findings offer significant insights into the regulation of BaGTA-mediated gene transfer in pathogens of the genus *Bartonella* and suggest a broader impact of processive antitermination as a regulatory principle in related GTAs.

## Results

### BrrG shows structural similarity to the P21 phage-encoded Q21 antiterminator and is essential for GTA transfer and expression of capsid components

Previously, we identified by multiplexed transposon sequencing the genes of BaGTA encoding both for donor transfer and recipient uptake functions (Québatte et al., 2017). However, many of the identified genes had only hypothetical or no functional sequence-based annotation, leaving their contribution to BaGTA function unclear. To address this knowledge gap, we employed the protein structure prediction tool AlphaFold2 to model unannotated proteins of BaGTA genes and searched the Protein Data Bank (PDB) database for similar structures. We discovered that the *brrG* gene (Fig. 1A), which is located in the *ror* locus, encodes a protein with structural similarity to the antiterminator Q21 of the lambdoid bacteriophage P21 (Fig. 1B). Q21 and its counterpart Q from phage λ (Qλ) function as processive antiterminators involved in bypassing intrinsic terminators (Guo et al., 1991; Grayhack and Roberts, 1982). This enables uninterrupted expression of the late genes, which encode capsid morphogenesis and lysis cassettes. Therefore, we hypothesized that, similar to Q21 and Qλ, BrrG represents a putative antiterminator controlling gene expression within the BaGTA regulatory network.

To test this hypothesis, we assessed the effect of *brrG* overexpression on the transcription of the *bgtA-K* locus that encodes components of the capsid synthesis and assembly machinery as essential components of BaGTA production. We utilized the previously established reporter strain *bgtC-gfp* that contains a promoter-less *gfpmut2* inserted in the *bgtA-K* locus in the intergenic region between *bgtC* and *bgtB* (Québatte et al., 2017) (Fig. 1A). This transcriptional reporter enabled us to monitor *bgtA-K* expression over time using flow cytometry (Fig. 1C, upper panel). We observed that plasmid-based overproduction of BrrG increased the fraction of GFP fluorescent bacteria up to 75%, compared to about 15% at native BrrG expression (Fig. 1D), thus massively expanding the fraction of cells expressing the *bgtA-K* locus. To test whether this increase is followed by GTA production and elevated DNA transfer, we performed the GTA transfer assay. For that, we cocultivated donor and recipient cells carrying different resistance markers followed by plating on selective media to enumerate double-resistant bacteria as a read-out for the frequency of GTA-mediated gene transfer (Fig. 1C, lower panel). We observed that - while the proportion of cells expressing the BaGTA locus increased as a result of BrrG overproduction, GTA transfer rates remained similar to native BrrG expression (Fig. 1E). In contrast, deleting *brrG* led to a significant decrease in the proportion of fluorescent bacteria (Fig. 1F) and caused DNA transfer rates to drop at least 100-fold compared to those observed at native BrrG expression (Fig. 1G). Together, these findings indicate that BrrG plays a crucial role in regulating BaGTA. It is capable of activating the expression of capsid components in a large part of the population that is initially GTA inactive. However, overproduction of BrrG is not sufficient to increase the rate of DNA transfer, indicating that it acts in concert with other regulators controling other crucial steps of this complex process.

### BrrG binds in proximity to the P*_bgtA-K_* promoter located upstream of to the putative *bgtA-K* operon

In lambdoid phages, both Q21 and Qλ are DNA-binding proteins that recognize a motif located in the promoter region of the late operon, which encodes lysis cassettes and proteins required for capsid morphogenesis (Guo et al., 1991; Grayhack and Roberts, 1982; Yin et al., 2019; Shi et al., 2019; Yin et al., 2022). Considering the structural similarity between BrrG and Q21, we hypothesized that BrrG may be functionally conserved and thus able to directly bind to DNA at the yet unidentified promoter of the *bgtA-K* locus. To test this, we performed chromatin immunoprecipitation of epitope-tagged BrrG followed by DNA sequencing. We found that BrrG binds to DNA, with a single binding site located in the intergenic region far upstream of the *bgtK* gene representing the first conserved gene of the *bgtA-K* locus (Fig. 2A, Suppl. Figure 2) (Guy et al., 2013; Tamarit et al., 2018). Given this remote upstream location of the BrrG binding site, we wondered whether the promoter elements might also be located nearby. First, we tested whether the 999 bp fragment spanning the region upstream of *bgtK* until the end of *clpB* could drive expression of a fused promoter-less *dsRed* gene using flow cytometry as readout (Fig. 2A, plasmid p*Pf3-dsRed*). Strikingly, this fragment did not only demonstrate promoter activity but also displayed at the population level a temporal control resembling closely the expression pattern of the chromosomal transcriptional reporter strain *bgtC-gfp H1^GmR^* with a characteristic peak in the size of expressing population at the 6 h time-point of the 24 h expression kinetic (Fig. 2B). This finding suggests that the *bgtA-K* locus is organized as a transcriptional unit (operon) that is driven by a promoter localizing to the 999 bp fragment upstream of *bgtK*, which we thus named P*_bgtA-K_* (Fig. 2A).

**Figure 2.**
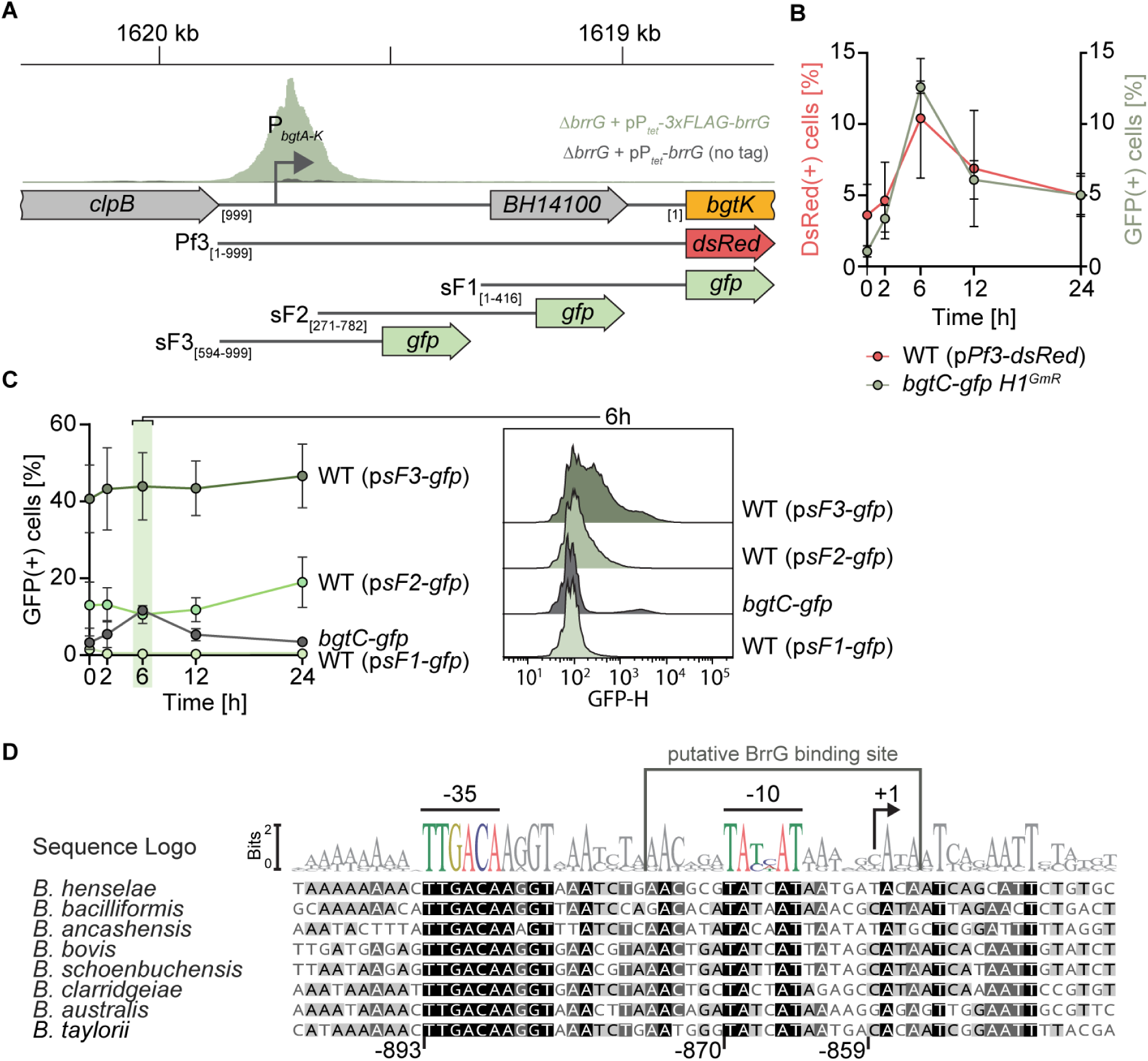
BrrG binds in proximity to the P*_bgtA-K_* promoter located upstream of the putative *bgtA-K* operon. **(A)** Location of the P*_bgtA-K_* promoter and of the BrrG binding site identified by ChIP-Seq (gray histogram). *ΔbrrG* + pP*_tet_-brrG* (no tag) is shown in dark grey; *ΔbrrG* + pP*_tet_-3x-FLAG*-*brrG* is shown in green. The fragments upstream of the *bgtA-K* locus were tested for promoter activity in promoter-probe plasmids as indicated. **(B)** The fusion of the 999 bp fragment spanning from the region between *bgtK* and *clpB* to *dsRed* (strain WT (P*_Pf3_-dsRed*) displays a temporal transcriptional profile similar to the chromosomal transcription reporter for the *bgtA-K* locus (strain *bgtC-gfp H1^GmR^* [*bgtC-gfp BH01490::Himar1^GmR^*]). **(C)** The P*_bgtA-K_* promoter is located far upstream of the *bgtA-K* locus. Left: Temporal transcriptional profiles controlled by fragments spanning nucleotide 1-416 (plasmid p*sF1*), nucleotides 271-782 (plasmid p*sF2*), or nucleotides 594-999 (plasmid p*sF3*) of the 999 bp fragment in Pf3. Right: Corresponding histograms of fluorescence distribution at the 6 h time-point. Strain *bgtC-gfp* displaying wild-type expression pattern of the *bgtA-K* locus was used as control. **(D)** DNA sequence alignment of the predicted promoter region and putative BrrG binding site within psF3 fragment with homologous sequences from various *Bartonella* species. The predicted -35 and -10 promoter elements and the inferred transcriptional start site (+1) are indicated. Numbers below indicate the location of the promoter elements relative to *bgtK*. (B, C) The time-course experiments were started with the incubation of CBA agar-grown bacteria in M199 medium supplemented with 10% FCS. The experiments were done in biological triplicates, error bars indicate SD.

Next, to narrow down the location of P*_bgtA-K_*, we split the 999 bp fragment into three partially overlapping fragments and tested them with the same approach using a *gfpmut2*-based promoter-probe plasmid (plasmids p*sF1*, p*sF2* and p*sF3,* Fig. 2A). We observed that the most upstream fragment spanning nucleotides 594-999 was able to drive broad expression of *gfpmut2* at population level (plasmid p*sF3* in Figure 2C). The fraction of fluorescent bacteria was substantially higher compared to the chromosomal transcriptional reporter strain *bgtC-gfp*, expanding beyond 40% at each time-point (Fig. 2C, Suppl. Figure 3). Importantly, the lack of temporal regulation as seen by the chromosomal transcriptional reporter strain *bgtC-gfp* and likewise the 999 bp fragment in plasmid Pf3 (compare Figs. 2B and 2C) suggests that absence of downstream control elements located in the more proximal region spanning nucleotides 1-593 converted P*_bgtA-K_* into a constitutive state. To map P*_bgtA-K_* based on conserved elements (-10 and -35 boxes) we aligned corresponding sequences from a selection of *Bartonella* species covering the genomic diversity of this genus. By this comparative approach, we identified a highly conserved sequence motive (TTGACA-N_17_-TATCAT) that shows almost full identity to the canonical sigma-70 promoter sequence of *E. coli* (-35 box: 100%, -10 box: >80%, Fig. 2D) (Campbell et al., 2002; Murakami et al., 2002). This suggests that P*_bgtA-K_*functions as a minimal promoter of the *bgtA-K* locus. Interestingly, the BrrG binding region coincides with P*_bgtA-K_*, suggesting that BrrG controls transcription either by direct binding to P*_bgtA-K_* or in the vicinity of it. Taken together, we identified P*_bgtA-K_* as a strong constitutive promoter located far upstream of the *bgtA-K* locus, driving its expression as a transcriptional unit along with more proximal elements required for controlling temporal expression at the population level. Additionally, we found that the putative antiterminator BrrG, similar to Q21 and Qλ, is a DNA-binding protein that directly binds to P*_bgtA-K_* and likely controls *bgtA-K* operon expression at population level.

### A long leader sequence attenuating expression is located downstream of the P*_bgtA-K_* promoter

To confirm that the identified conserved sequence is the promoter and to hint at the function of the long downstream sequence in transcription, we generated a panel of promoter-probe plasmids harbouring truncations of the 999 bp fragment bearing promoter activity (p*F0-gfp* to p*F6-gfp*) (Fig. 3A). We observed that the 29 bp fragment spanning the predicted P*_bgtA-K_* displayed fluorescence in the vast majority of cells, confirming the promoter prediction (F0 fragment, Fig. 3B). Similar promoter activities were recorded for the fragments F1 and F2 that are extended by 27 bp or 133 bp towards the *bgtA-K* locus, respectively. A striking difference in fluorescence distribution, with an almost complete loss of fluorescent bacteria, was observed for the F3 fragment, which contains an additional 139 bp relative to F2 (Fig. 3B, Suppl. Figure 4). Fragments F4-F6 with further downstream extensions also displayed pattern of low fluorescence, while a bi-model distribution with low- and intermediate-expressing bacteria was observed for fragments F4 and F6.

**Figure 3.**
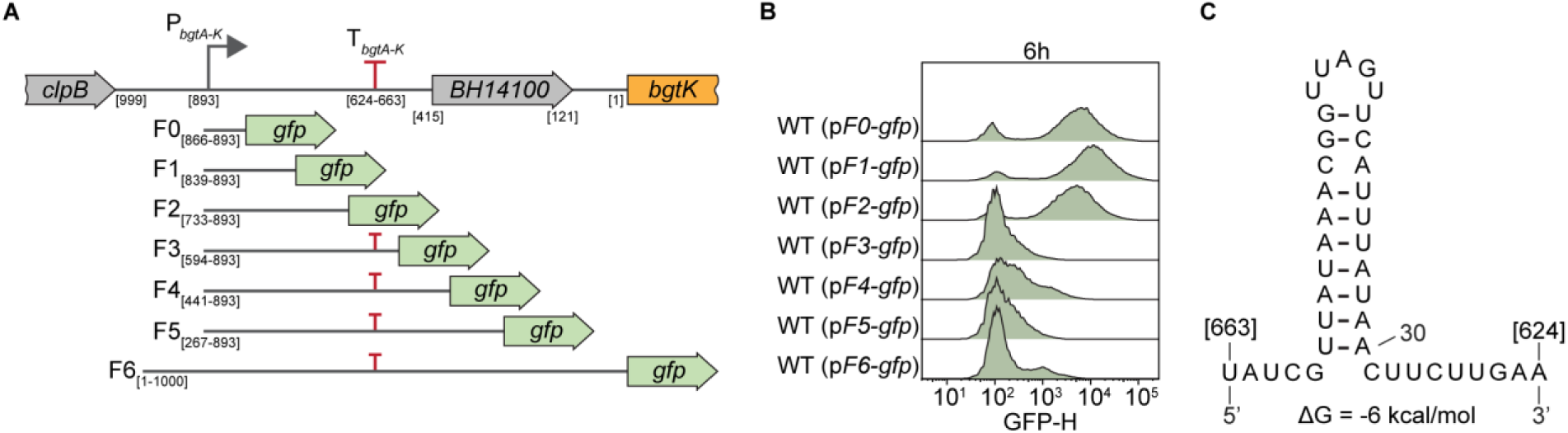
A putative terminator is encoded in the long non-protein coding sequence downstream of the P*_bgtA-K_* promoter. **(A)** Schematic representation of the 5’-part of the *bgtA-K* locus with the truncations tested with promoter-probe plasmids. Locations of the P*_bgtA-K_*promoter and the predicted terminator are shown. **(B)** Histograms of fluorescence distribution measured for the bacterial populations containing the promoter-probe plasmids with various truncations at 6 h. **(C)** Predicted secondary structure and free energy of the putative T*_bgtA-K_* terminator. Numbers in parentheses indicate the location of the terminator relative to *bgtK*. The time-course experiment was started with the incubation of CBA plate-grown bacteria in M199 medium supplemented with 10% FCS. Experiments were performed in biological triplicates.

Because fragment F3 is the shortest fragment displaying reduced expression, we hypothesized that a terminator could be present in the sequence extending the highly fluorescent F2 fragment. Indeed, a putative L-shaped terminator with a free energy (ΔG) of -6 kcal/mol is predicted within the stretch of 139 bp of the F3 fragment, which might explain the observed difference in expression levels (Fig. 3C). Altogether, these results demonstrate that the minimal promoter enables uniform and constitutive expression of the BaGTA locus at population level. The observed temporal regulation and heterogeneity of the gene cluster expression appear to be provided by the stretch of 864 bp located downstream of the promoter and upstream to *bgtK* as first gene of the *bgtA-G* locus. This stretch of DNA contains the putative terminator and could harbour other terminators or additional regulatory elements.

### A -10-like element is required for expression of the *bgtA-K* locus

In phage lambda, the process of Q-mediated antitermination relies on the engagement of Q into the complex with RNA-polymerase (RNAP) and the conversion of the latter into a termination-resistant form (Perdue and Roberts, 2011). This process begins with RNAP pausing at the -10- like element, a six-nucleotide motif (NANNNT), located just downstream of the -10 promoter element, which allows Q to engage with the transcription complex (Perdue and Roberts, 2011).

Therefore, we set out to determine whether a -10-like element is present in the vicinity of the identified P*_bgtA-K_* minimal promoter. We examined the previously generated alignment and identified a conserved NANNNT site located 5 bp downstream of the identified promoter P*_bgtA-K_*(Fig. 4A). As the conserved nucleotides A and T at positions +2 and +6 in phage lambda and other phages are also conserved in this motif directly downstream of P*_bgtA-K_*, we hypothesized that this site could act as a -10-like element (Fig. 4A). To validate this hypothesis, we used the promoter-probe plasmid containing the F6 fragment (Fig. 3A), and introduced two point mutations, replacing the conserved A in position +2 to G and T in position +6 to C (p*F6_(-10mut)_-gfp*), respectively. Additionally, we introduced a second plasmid directing aTc-inducible BrrG overproduction to test whether these mutations would affect the activity of the putative antiterminator in the flow cytometry assay (Fig. 1C). In contrast to plasmid p*F6-gfp* encoding the native -10-like element (Fig. 4B, left, -aTc), p*F6_(-10mut)_-gfp* did not display the characteristic kinetics of GFP-expression of wild-type bacteria with a peak at 6 h (Fig. 4B, right, -aTc, Suppl. Figure 5). Moreover, neither did it display the massive expansion of the GFP-expressing population upon BrrG overproduction (+aTc) (Fig. 4B, Suppl. Figure 5). Thus, the replacement of two conserved nucleotides at the putative -10-like element completely abolished transcription of the downstream *bgtA-K* locus under both native BrrG expression and BrrG overproduction conditions. This suggests that the identified conserved site is a functional -10-like element essential for the BrrG-mediated antitermination.

**Figure 4.**
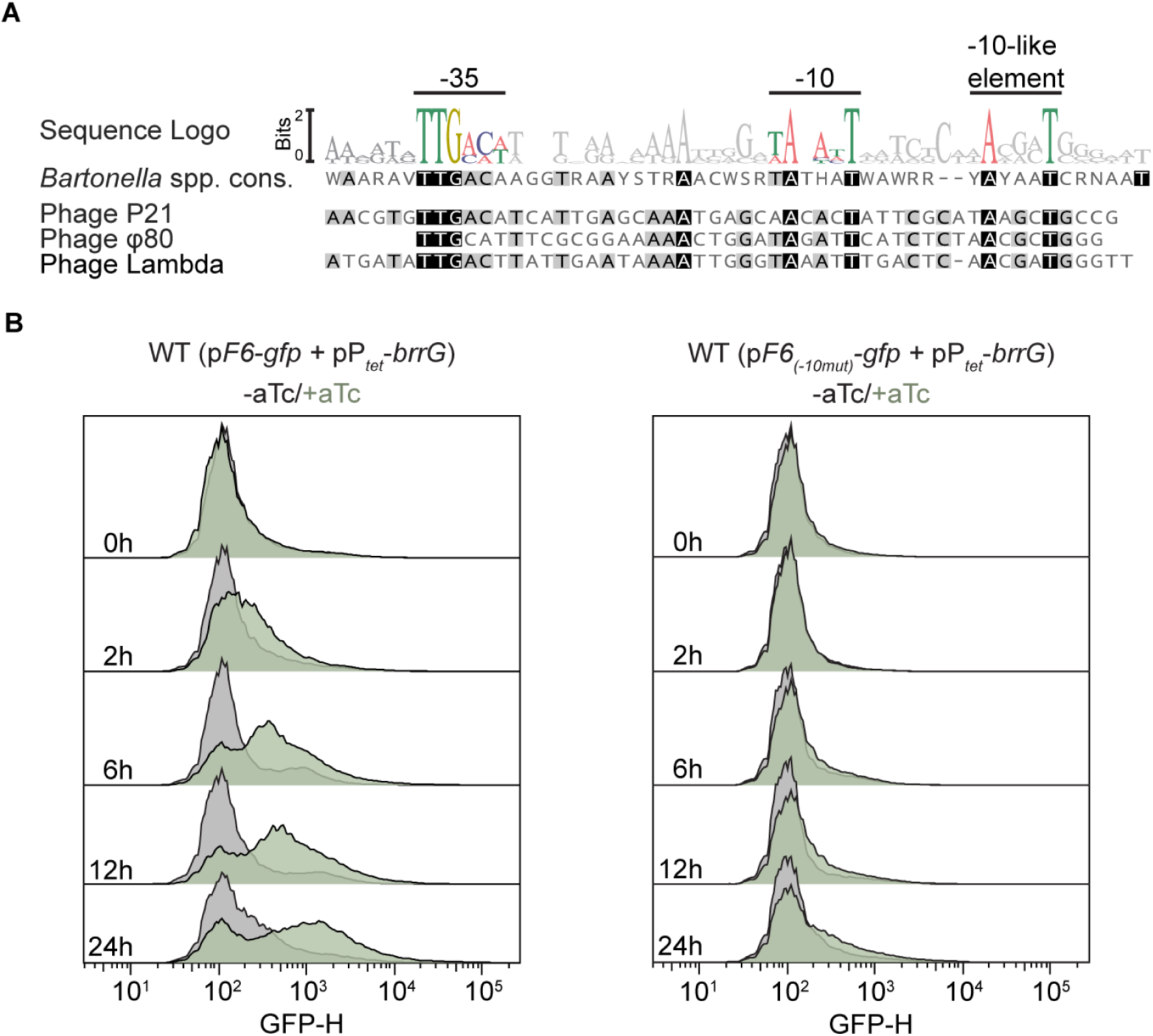
The -10-like element downstream of the P*_bgtA-K_* promoter is crucial for the antitermination by BrrG. **(A)** Alignment of *Bartonella* spp. consensus and lambdoid phage sequences (P21, phi80 and lambda) sequences containing P*_bgtA-K_* and the P*_R’_* promoters followed by the -10-like elements, respectively. Promoter -35 and -10 elements, as well as the -10-like elements are marked. **(B)** Histograms of fluorescence distribution for the bacterial populations containing the promoter-probe plasmid pF6 (compare to Fig. 3A) with the P*_bgtA-K_*promoter and the adjacent native -10-like element or its derivative, plasmid p*F6_(-10mut)_*, carrying mutations in two conserved positions of the -10-like element. Plasmid pP*_tet_-brrG* encodes an aTc-inducible *brrG* expression cassette. BrrG overproduction was induced by 25 ng/ml aTc. The time-course experiment was started with the incubation of CBA plate-grown bacteria in M199 medium supplemented with 10% FCS. Inducer aTc was added at time-point 0 h. Experiments were performed in biological triplicates.

### BrrG is a processive antiterminator allowing RNAP to bypass the termination site

We showed that BrrG shares structural features with the processive antiterminator Q21 and that its function is dependent on the -10-like element suggesting that BrrG belongs to the class of processive antiterminators. To validate this hypothesis, we tested the antitermination properties of BrrG towards a synthetic terminator. For this, we used the artificially designed strong terminator BBa_B1006 from the iGEM collection and placed it in between the F2 fragment (see Fig. 3A), which contains the native P*_bgtA-K_* promoter and downstream -10-like element, but not the native terminator and the *gfp* reporter gene (Fig. 5A). The plasmid p*F2-gfp* containing the terminator-less fragment F2 drives GFP-expression in a large part of the bacterial population at both BrrG native expression and overproduction conditions (Figs. 3B and 5B). In contrast, the derivative p*F2-T_BBa_1006_-gfp* containing in addition the artificial terminator showed a strongly reduced population expression of GFP, but a marked increase in GFP-expressing bacteria upon BrrG overproduction (Fig. 5B, right panel, Suppl. Figure 6). Together, these findings demonstrate that the presence of BrrG allows the RNAP to bypass both the native termination site, as well as an unrelated synthetic terminator, indicating that BrrG is indeed a processive antiterminator.

**Figure 5.**
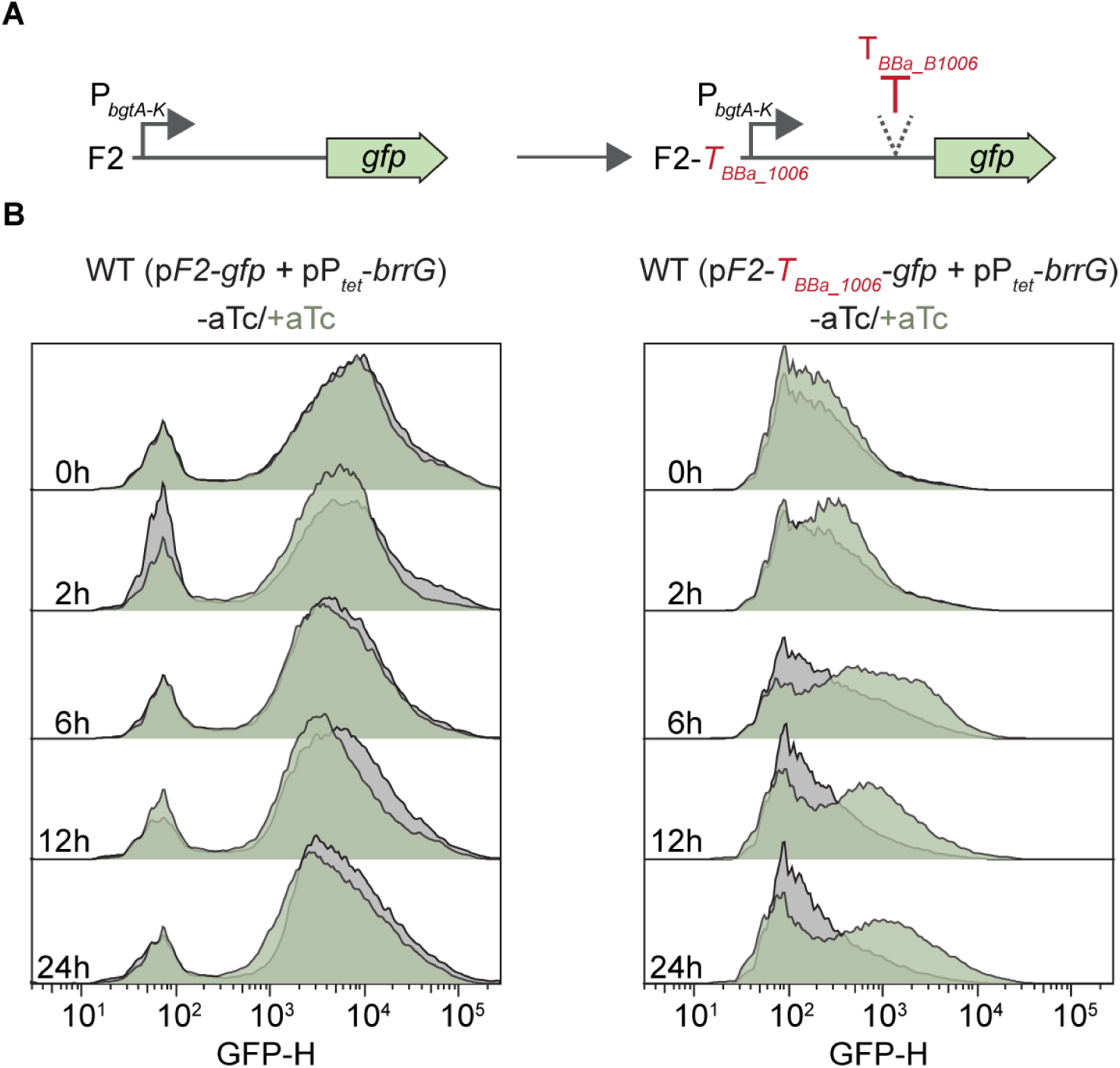
Antiterminator BrrG allows RNA polymerase to bypass an engineered termination site. **(A)** Schematic representation of the experimental setup. Plasmid p*F2-T_BBa_1006_-gfp* carries the insertion of the artificial strong terminator *T_BBa_1006_* in between the terminator-less promoter fragment F2 and *gfp* of plasmid pF2-*gfp.* **(B)** Histograms of fluorescence distribution for the bacterial populations containing the fragment F2, the fragment F2 with the terminator and the sequence with the mutated -10-like element (pF6_-10mut_). Time of the measurements is indicated. Induction was done with 25 ng/ml aTc. Bacteria were cultivated in M199 medium supplemented with 10% FCS and aTc inducer at indicated concentrations. Experiments were performed in biological triplicates.

**Figure 6.**
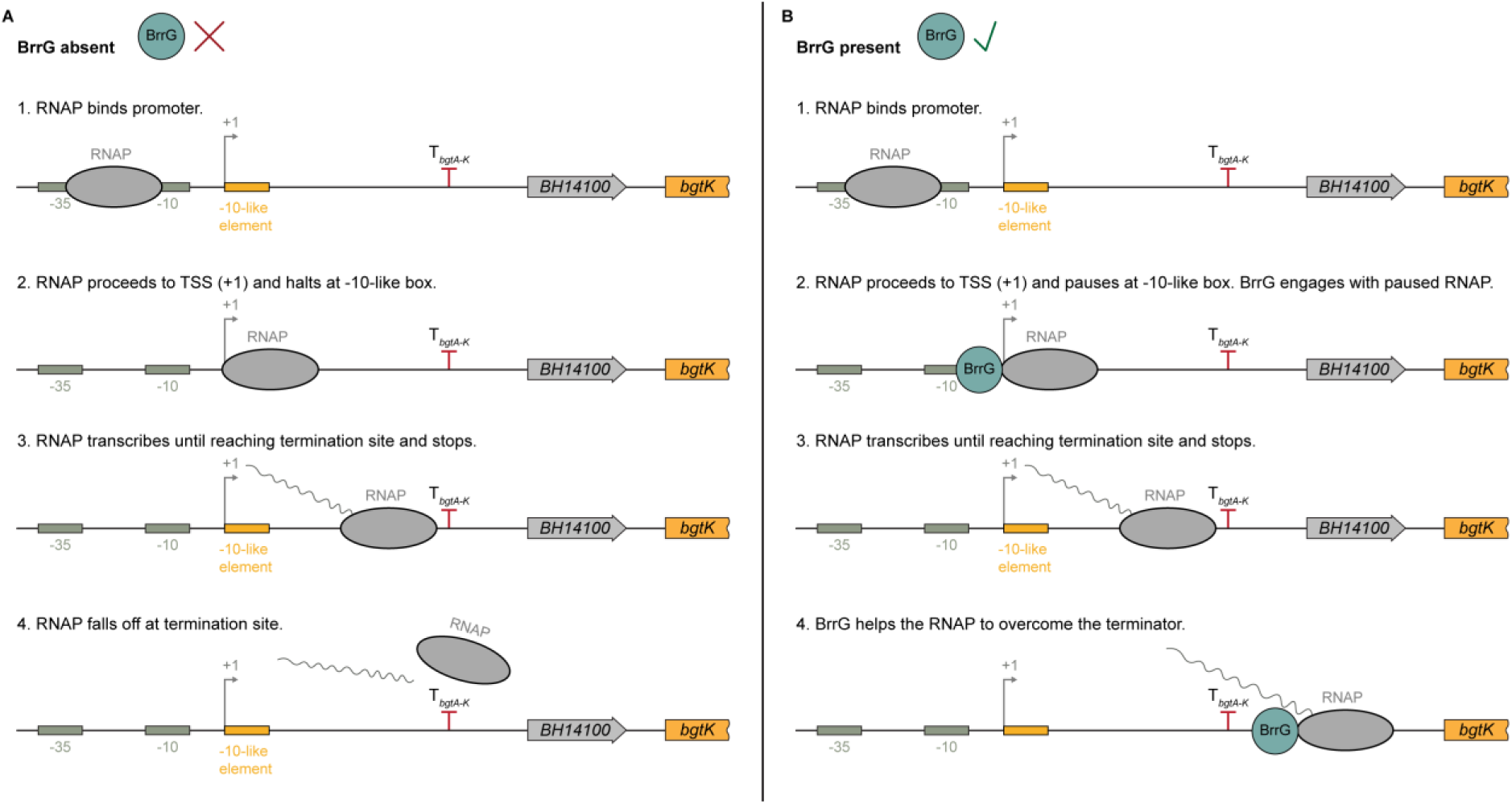
Model of BrrG-antitermination. **(A)** Schematic representation of terminator-dependent transcription inhibition. RNAP initiates transcription at the TSS (+1) and proceeds until the termination site (T*_bgtA-K_*), where it falls off. **(B)** Transcriptional antitermination of the BaGTA in presence of BrrG. BrrG acts as antiterminator enabling the transcription of the downstream *bgtA-K* cassette.

## Discussion

Processive antitermination is an important mechanism of transcriptional control identified both for bacteria and phages. In temperate phages processive antiterminators, i.e. phage lambda N and Q-antiterminators, switch on expression of early and late genes, respectively, and thus enable the progression of the infection cycle (Goodson and Winkler, 2018).

Given the ancestral relationship between GTAs and phages, this study investigates whether transcription attenuation and antitermination are preserved in the chimeric BaGTA and whether they play a role in the regulatory integration of two functional gene clusters of divergent phage origin. We showed that the “late” capsid producing *bgtA-K* genes belong to a single transcriptional unit derived from one phage, and that their expression is enabled by the antiterminator BrrG, encoded within the *ror* locus, which originates from a second phage. However, the question remains: Is BrrG a Q-like antiterminator, and does it function in a similar manner?

The canonical Q-dependent regulatory circuit consists of several critical components: Q antiterminator, the promoter P*_R’_*, a termination site and a -10-like element (Perdue and Roberts, 2011; reviewed in Casjens and Hendrix, 2015). The first component is the Q antiterminator which binds to DNA and engages into the complex with RNAP. The Q-RNAP complex is termination-resistant and permits RNAP to bypass the termination signals in front of late genes which results in the synthesis of the capsid components (Guo et al., 1991; Grayhack and Roberts, 1982). In our study we showed that BrrG has structural similarity to the Q21 antiterminator of the P21 lambdoid phage. Moreover, BrrG is able to counteract transcription attenuation imposed by an introduced artificial termination site. Additionally, we demonstrated that BrrG binds in proximity to the P*_bgtA-K_* promoter and upregulates transcription of the *bgtA-K* cluster – BaGTA’s counterpart of the late gene cluster.

The second component of the canonical Q-dependent regulatory circuit is the promoter P*_R’_* containing Q-binding site for interaction with Q and engagement with RNAP. In this study, we demonstrated that the P*_bgtA-K_* promoter overlaps with the BrrG binding site. Given that the promoter P*_bgtA-K_* is located 864 bp upstream from the first gene of the *bgtA-K* cassette and is separated from the former by a gene BH14100, this raises the question of whether BH14100 belongs to the *bgtA-K* cluster and what functional role it may play. Previous studies have shown that BH14100 is not conserved among *Bartonellae* and thus is likely not involved in the BaGTA regulation or synthesis (Guy et al., 2013; Tamarit et al., 2018). In support of this, our previous study using multiplexed transposon sequencing showed no effect of BH14100 disruption on GTA transfer, indicating that BH14100 has either no or only a very modest effect on GTA synthesis (Québatte et al., 2017).

The third component of the canonical Q-dependent regulatory circuit is the termination site which attenuates expression of the late gene cluster_’_. We showed that the putative terminator T*_bgtA-K_*is located downstream of the P*_bgtA-K_* promoter and regulates expression of the *bgtA-K* operon. Whereas the *bgtA-K* locus is only active in the GTA producing cells, deleting the putative terminator T*_bgtA-K_*renders broad and constitutive expression of the *bgtA-K* locus which suggests that the temporal regulation and heterogeneity in expression are granted by T*_bgtA-K_*.

The fourth component of the canonical Q-dependent regulatory circuit is the -10-like element located in proximity to the promoter P_R’_ driving expression of late genes. This element resembles the -10-element of the promoter and defines the sigma-dependent pausing of RNAP which is crucial for the engagement of Q into the complex with RNAP (Perdue and Roberts, 2011). We found that expression of the *bgtA-K* locus is dependent on the conserved -10-like element located downstream of the P*_bgtA-K_* promoter. Altogether, these data suggest that BrrG is an antiterminator found in BaGTA and involved in regulating expression of the *bgtA-K* cluster. Based on these lines of evidence we propose BrrG as Q-like processive antiterminator.

However, why does GTA-mediated DNA transfer remain unaffected upon *brrG* overexpression?

A simple explanation could be that bacterial lysis and particle release are controlled independently of BrrG which activates only transcription of the *bgtA-K* locus. This notion is supported by the apparent absence of lysis of fluorescent bacteria upon BrrG overproduction (Fig. 1D). For the archetypical RcGTA of *Rhodobacter capsulatus*, particle synthesis and release are controlled differentially by the CckA-ChpT-CtrA phosphorelay system, where the phosphorylation state of CtrA controls the progression from the GTA synthesis to a lytic release (Farrera-Calderon et al., 2021; Fogg, 2019; Westbye et al., 2018; Westbye et al., 2013). We hypothesize that similarly to RcGTA a trigger could be involved in the regulation of bacterial lysis and BaGTA release. BaGTA contains two putative lysis cassettes – *bgtAB* and *bgtTUV* – and only the former one is located within the larger *bgt* cluster, suggesting that the latter cassette is controlled independently.

Taken together, we demonstrate that the transcriptional control mechanism involving transcription attenuation and antitermination, as described for phage lambda, is functionally conserved in the chimeric BaGTA. This mechanism specifically coordinates the temporal expression of the ROR and *bgtA-K* loci, which have independent phage origins. The ROR-encoded BrrG acts as a processive antiterminator, facilitating the induction of the *bgtA-K* transcriptional unit. However, is this regulatory mechanism also conserved in other types of GTA beyond BaGTA of *Bartonella*? The recent publication by Tran and Le has shown that termination-antitermination is involved in control of gene expression in evolutionary distinct CcGTA of *Caulobacter crescentus* (Tran and Le, 2024). The authors showed that transcription of the structural genes, which are counterparts of the *bgtA-K* locus, is co-activated by GafY and IHF, while the GafZ antiterminator prevents expression attenuation at the termination sites. Furthermore, GafYZ regulators of CcGTA are homologous to N’- and C’-domains of the highly conserved GafA – the direct activator of RcGTA (Fogg, 2019; Gozzi et al., 2022; Sherlock and Fogg, 2022), which indicates that RcGTA might also be controlled by processive antitermination. Previous studies of GafA indicated that its C’-domain has similarity to the region 4 of a sigma factor (R4σ) and therefore authors proposed GafA as an alternative sigma factor or a transcription factor (Fogg, 2019; Sherlock and Fogg, 2022). However, the related R4σ-like fold was also found in the resolved structure of Q21 antiterminator indicating that findings of Fogg and Sherlock might, in fact, speak for the implication of GafA in antitermination (Shi et al., 2019). Interestingly, Tran and Le found neither sequence nor structural similarity of GafZ to known antiterminators Qλ and Q21. Our comparison of resolved structure of Q21 and modelled BrrG, GafZ and GafA, clearly indicates similar structural features, suggesting both GafZ and GafA being antiterminators (Suppl. Figure 1).

RcGTA and CcGTA, on one hand, and BaGTA on the other, belong to two evolutionary distinct groups of GTA that emerged from independent events of phage domestication (Kogay et al., 2022; Tamarit et al., 2018). The reliance of these groups on the termination-antitermination mechanism of transcription control clearly suggests that processive antitermination is a common regulatory mechanism in GTAs, which evolved independently in two types of GTA within Alphaproteobacteria. To date, five types of GTAs have been identified in both bacteria and archaea. Given their phage heritage, it remains unclear whether other GTA types also depend on processive antitermination for regulation, and this should be explored in future studies.

## Materials and Methods

### Bacterial Strains and growth conditions

All bacterial strains, plasmids and oligonucleotides used in this study are listed in Supplementary Tables 1, 2 and 3, respectively. *B. henselae* strains were grown at 35°C on Columbia blood agar (CBA) supplemented with 5% sheep blood, appropriate antibiotics (gentamicin (Gm) – 10 μg/ml, kanamycin (Km) – 30 μg/ml, streptomycin (Sm) – 100 μg/ml, spectinomycin (Sp) – 50 μg/ml) and other supplements (isopropyl β-D-thiogalactosidase (IPTG) – 500 μM, anhydrotetracycline (aTc) – 10 ng/ml or 25 ng/ml) in a humidified atmosphere containing 5% CO_2_. *B. henselae* strains were stored as cryostocks at -80°C and prior the experiment seeded on CBA for 3 days and subsequently expanded and grown on fresh plates for 2 days. *E. coli* strains were stored as cryostocks at -80°C and were grown at 37°C overnight in lysogeny broth (LB) or on solid LB agar plates (LA) supplemented with appropriate antibiotics (ampicillin (Ap) – 100 μg/ml, Gm – 10 μg/ml, Km – 30 μg/ml) and other supplements (2,6-diaminopimelic acid (DAP) – 1 mM, 1% glucose).

### Construction of strains

All bacterial strains used in this study are listed in Table 1.

**Table 1.**
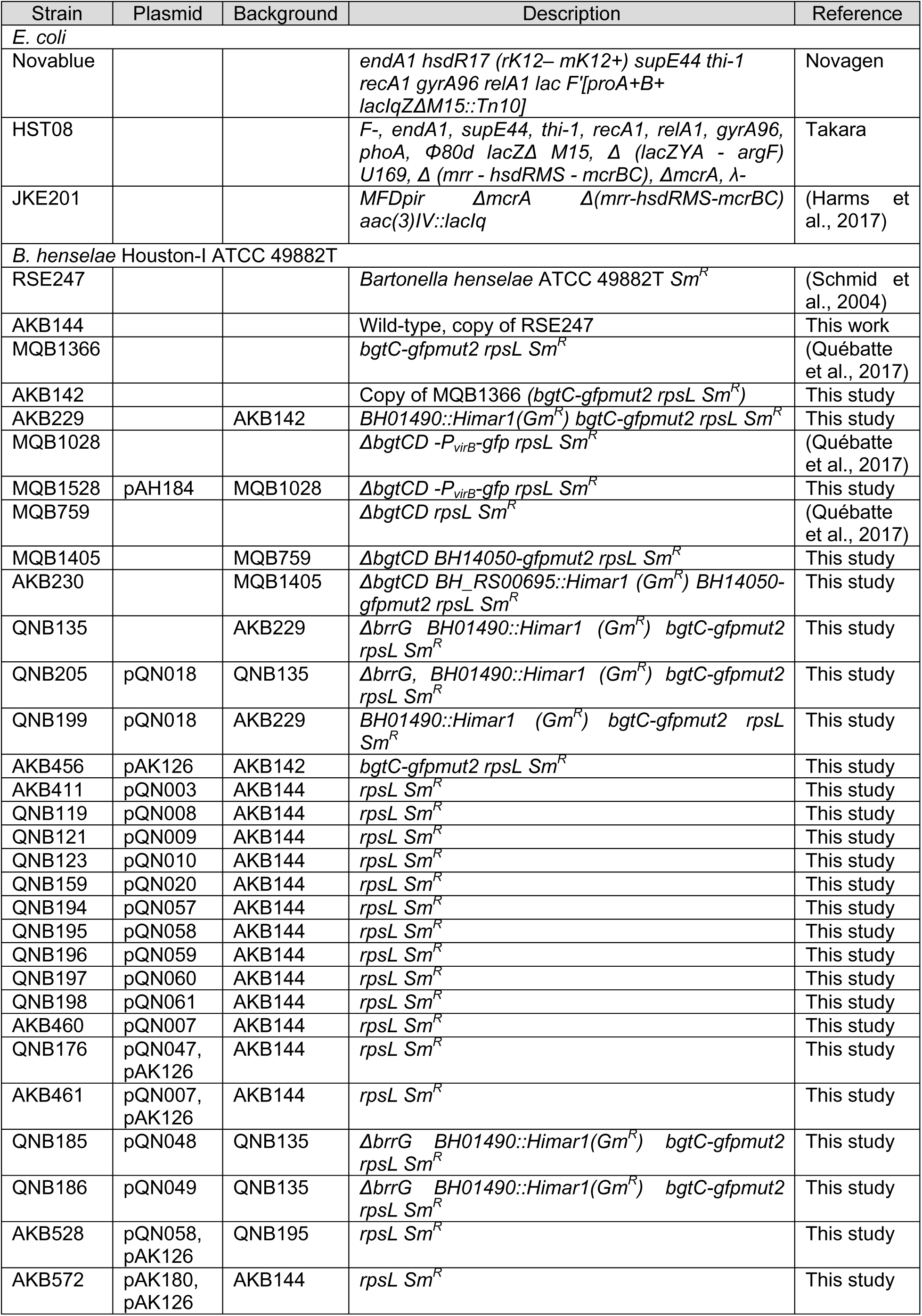
List of strains used in this study.

DNA manipulations were performed according to standard techniques and all cloned inserts were DNA sequenced to confirm sequence integrity. Chromosomal insertions or deletions in *B. henselae* were generated by a two-step gene replacement procedure as described in Schulein and Dehio (Schulein and Dehio, 2002). Plasmids were introduced into *B. henselae* strains by conjugation with *E. coli* strain JKE201 (MFD*pir*) using two-parental mating as described previously (Harms et al., 2017).

### Construction of plasmids

All plasmids used in this study are listed in Table 2 and all oligonucleotides used in this study are listed in Table 3. Plasmids generated in this study were constructed as follows:

pQN011: Plasmid pTR1000 was digested with the restriction enzyme BamHI. Homologous fragments 1 (upstream) and 2 (downstream) were PCR amplified from heat-killed *B. henselae* cells with primers prQN029 and prQN030, prQN031 and prQN032, respectively.

Those two fragments were cloned together by performing overlap extension PCR with primers prQN030 and prQN032. The fragment and the backbone were fused with the 5X In-Fusion® HD Enzyme Premix kit.

pQN018: *brrG* was PCR amplified from heat-killed *B. henselae* cells with primers prQN053 and prQN054. *tetR* was PCR amplified with primer prAK227 and prAK208 from 705-bp *tetR-P_tet_* promoter template ordered as a synthetic fragment from Genescript. The PCR product generated by amplifying pBZ485_a_empty with prAK089 and prAK109 was used as a backbone. Three fragments were combined with the 5X In-Fusion® HD Enzyme Premix kit.

pAK126: *brrG* was PCR amplified from heat-killed *B. henselae* cells with primers prAK440 and prAK441. *tetR* was PCR amplified with primer prAK444 and prAK208 from 705-bp *tetR-P_tet_* promoter template ordered as a synthetic fragment from Genescript. The PCR product generated by amplification of pPG100 with prAK116 and prAK167 was used as a backbone. Three fragments were combined with the 5X In-Fusion® HD Enzyme Premix kit.

pQN003: Fragment Pf3 was PCR amplified from heat-killed *B. henselae* cells with primers prQN005 and prQN014. *dsRed* was PCR amplified from pJC44 with primer prAK097 and prAK110. The PCR product generated by amplifying pBZ485_a_empty with prAK089 and prAK109 was used as a backbone. Three fragments were combined with the 5X In-Fusion® HD Enzyme Premix kit.

pQN006: g*fpmut2* was PCR amplified from plasmid pCD353 with primers prQN015 and prQN016. The PCR product generated by amplifying pBZ485_a_empty with prAK089 and prAK109 was used as a backbone. Two fragments were combined with the 5X In-Fusion® HD Enzyme Premix kit.

pQN008: Plasmid pQN006 was digested with BamHI. Fragment sF1 was PCR amplified form heat-killed *B. henselae* cells with primers prQN021 and prQN022. The fragment and the backbone were ligated with the 5X In-Fusion® HD Enzyme Premix kit.

pQN009: Plasmid pQN006 was digested with BamHI. Fragment sF2 was PCR amplified form heat-killed *B. henselae* cells with primers prQN018 and prQN017. The fragment and the backbone were ligated with the 5X In-Fusion® HD Enzyme Premix kit.

pQN010: Plasmid pQN006 was digested with BamHI. Fragment sF3 was PCR amplified form heat-killed *B. henselae* cells with primers prQN019 and prQN020. The fragment and the backbone were ligated with the 5X In-Fusion® HD Enzyme Premix kit.

pQN020: The minimal predicted P_GTA_ promoter was introduced into plasmid pQN006 by PCR with the primers prQN061 and prQN062. The plasmid was circularized with the 5X In-Fusion® HD Enzyme Premix kit.

pQN057: The PCR product generated by amplifying pQN006 with prQN111 and prAK109 was used as a backbone. F1 was PCR amplified from heat-killed *B. henselae* cells with primers prQ134 and prQN135. F1 and the backbone were combined with the 5X In-Fusion® HD Enzyme Premix kit.

pQN058: The PCR product generated by amplifying pQN006 with prQN111 and prAK109 was used as a backbone. F2 was PCR amplified from heat-killed *B. henselae* cells with primers prQ134 and prQN136. F2 and the backbone were ligated with the 5X In-Fusion® HD Enzyme Premix kit.

pQN059: The PCR product generated by amplifying pQN006 with prQN111 and prAK109 was used as a backbone. F3 was PCR amplified from heat-killed *B. henselae* cells with primers prQ134 and prQN137. F3 and the backbone were ligated with the 5X In-Fusion® HD Enzyme Premix kit.

pQN060: The PCR product generated by amplifying pQN006 with prQN111 and prAK109 was used as a backbone. F4 was PCR amplified from heat-killed *B. henselae* cells with primers prQ134 and prQN138. F4 and the backbone were ligated with the 5X In-Fusion® HD Enzyme Premix kit.

pQN061: The PCR product generated by amplifying pQN006 with prQN111 and prAK109 was used as a backbone. F5 was PCR amplified from heat-killed *B. henselae* cells with primers prQ134 and prQN139. F5 and the backbone were ligated with the 5X In-Fusion® HD Enzyme Premix kit.

pQN007: Fragment F6 was PCR amplified from heat-killed *B. henselae* cells with primers prQN005 and prQN014. Plasmid pQN006 and F6 were restricted with BamHI and Xmal and ligated with RAPID DNA Dephos and Ligation Kit.

pQN047: The mutated -10-like sequence was introduced into plasmid pQN007 by PCR with primers prQN120 and prQN121. The plasmid was circularized with the 5X In-Fusion® HD Enzyme Premix kit.

pQN048: N’-3x-FLAG was introduced into pQN018 by PCR with primers prQN114 and prQN115. The plasmid was circularized with the 5X In-Fusion® HD Enzyme Premix kit.

pQN049: C’-3x-FLAG was introduced into pQN018 by PCR with primers prQN116 and prQN117. The plasmid was circularized with the 5X In-Fusion® HD Enzyme Premix kit.

pAK180: Terminator T*_BBa_1006_* was introduced into pQN058 by PCR with primers prAK592 and prAK593. The linear PCR product was used for the direct transformation of *E. coli*.

**Table 2.**
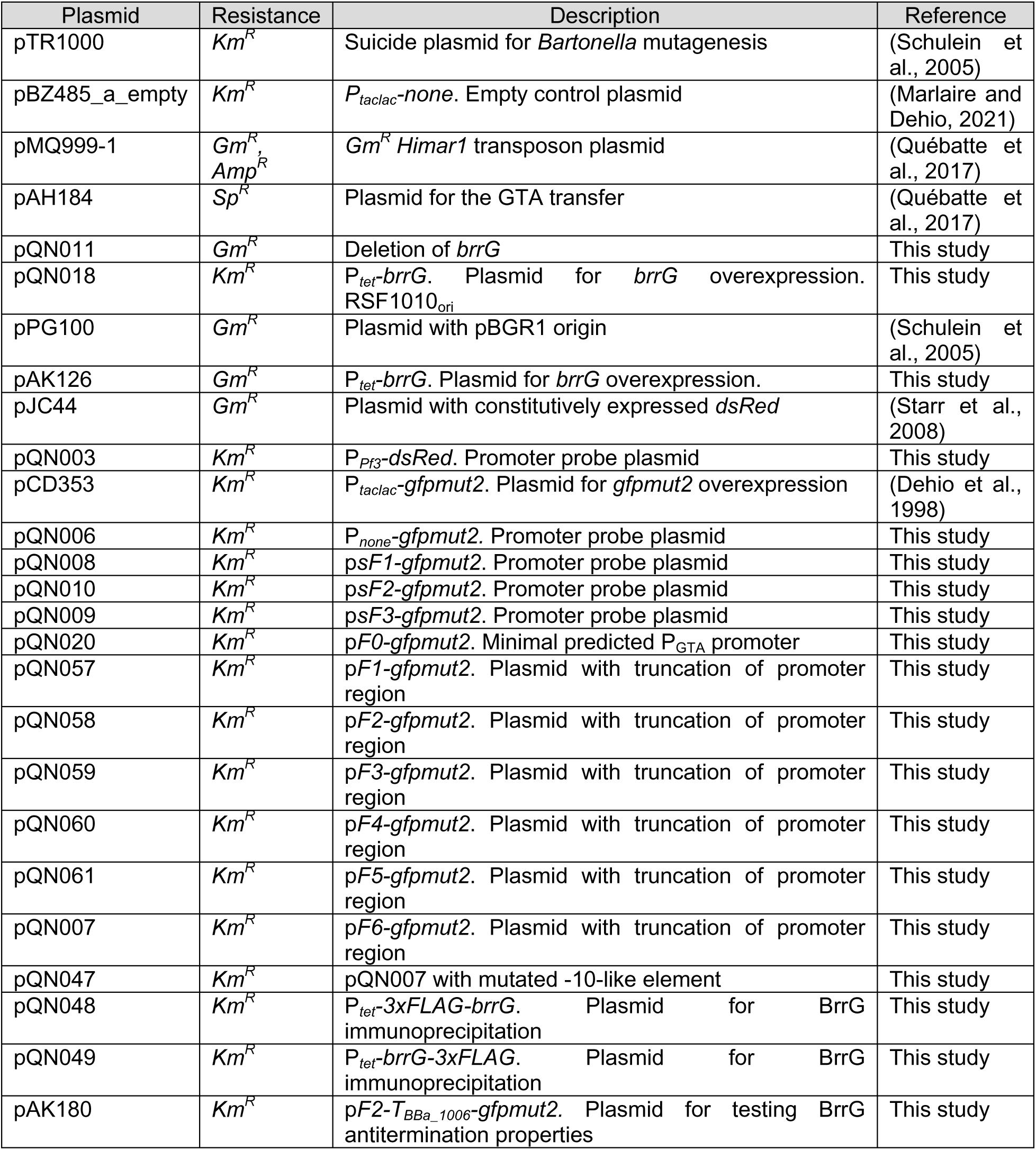
List of plasmids used in this study.

**Table 3.**
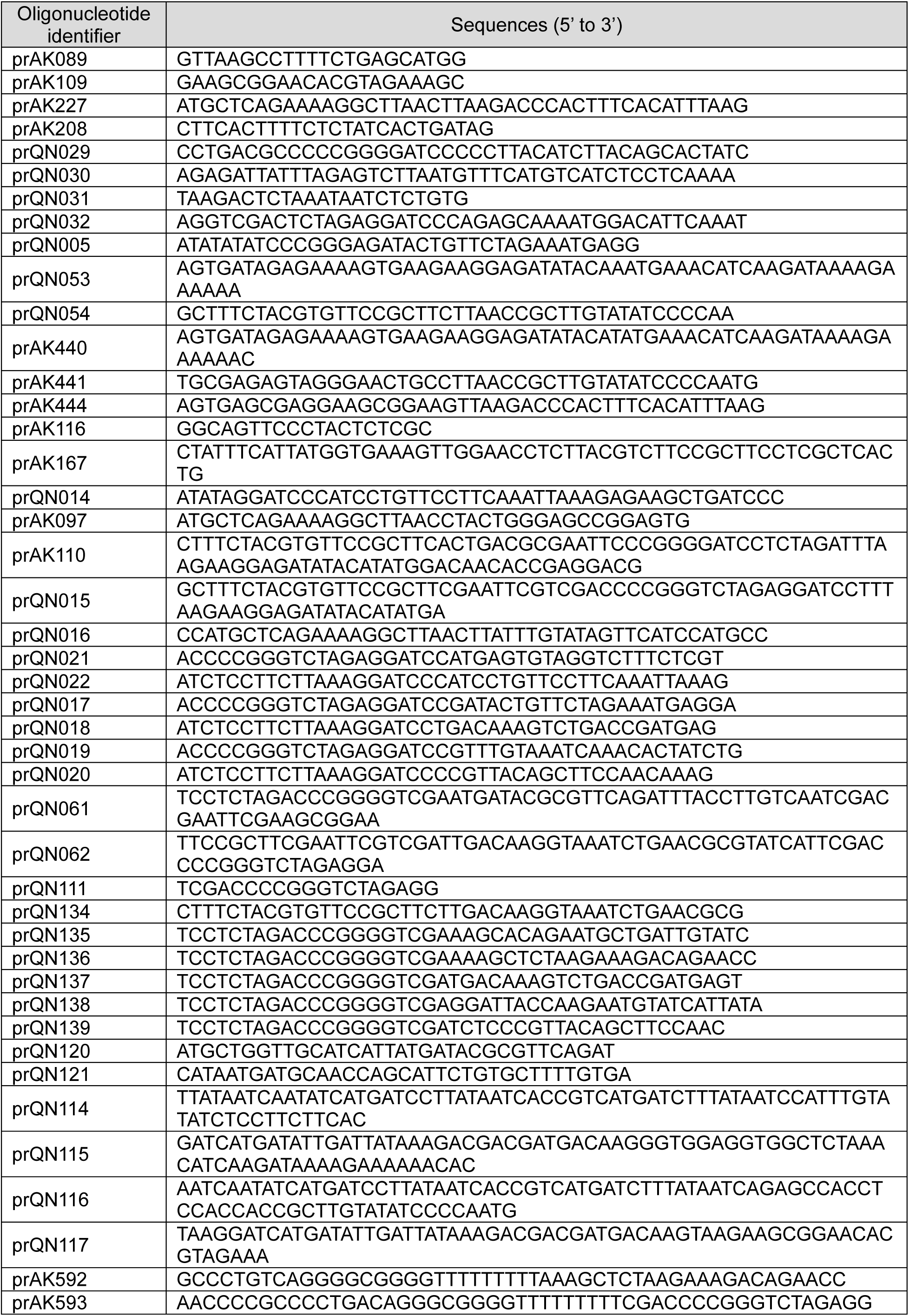
List of oligonucleotides used in this study.

### Measuring expression of the BaGTA locus with flow cytometry

*B. henselae* strains were grown as described above. Bacteria were resuspended in M199 supplemented with 10% FCS and other additives, e.g. inducer, if required (hereafter M199+), diluted to OD_600 nm_ = 0.008 and incubated in 24-well plate for a defined amount of time in a humidified atmosphere at 35°C and 5% CO_2_. After a defined amount of time bacteria were collected, centrifuged, and washed once with PBS. Bacteria were resuspended in 4% PFA and fixed for 7 min at RT. The fixation was stopped by adding PBS supplemented with 2% FCS. Samples were kept at 4°C in the dark before flow cytometry analysis. GFP fluorescence in samples was recorded with the BD LSRFortessa™ Cell Analyzer at the FACS Core Facility, Biozentrum, the University of Basel. Data were analysed using software FlowJo (Version 10.8.1). Visualisation of the data was done with GraphPad prism 8.

### GTA transfer assay

GTA transfer assays were done as co-cultivations of two *B. henselae* strains. *B. henselae* strains were grown as described above. Bacteria were resuspended in M199+ supplemented, if required, with inducer at specified concentrations and diluted to OD_600 nm_ = 0.008. For tested co-cultivation pair 1 ml of each strain was mixed and a mixture was incubated in 24-well plate for 24 h in a humidified atmosphere at 35°C and 5% CO_2_. After 24 h bacteria were collected and centrifuged. Bacterial pellets were resuspended in 30 μl of M199+ and resuspensions were used for serial dilutions followed by plating onto CBA plates supplemented with SmGm or SmGmSp antibiotics. Bacteria were grown for 6-15 d in a humidified atmosphere at 35°C and 5% CO_2_. Bacterial colonies were counted, and CFU/ml was determined for each co-cultivation pair. Frequency of double resistant bacteria (DRB) was calculated as the ratio of CFUs of recipient DRB (Sm^R^ Gm^R^ Sp^R^) / total number of recipients (Sm^R^ Gm^R^), yielding the transfer frequency per recipient cell.

### Chromatin immunoprecipitation coupled with high throughput sequencing

*B. henselae* strains were grown as described above. Bacteria were harvested from plates and resuspended in 5 ml of Schneider’s Insect Medium (24.5 g/l Schneider’s Drosophila Powder Medium Revised, 5% (wt/vol) sucrose, 0.025 M HEPES, adjusted pH 7.25) supplemented with 10% FCS (hereafter Schneider’s Medium) (Guy et al., 2013; Riess et al., 2008). Bacteria were diluted to OD_600 nm_ = 0.01 in 500 ml Schneider’s Medium for each strain and grown at 35°C with shaking 135 rpm for 4 d until OD_600 nm_ = 0.6-0.7. Bacteria were pelleted and pellets were resuspended in 50 ml PBS. Bacteria were fixed with PFA with final concentration 1% and incubated for 10 min at RT. The fixation was stopped by adding glycine to a final concentration of 125 mM, and mixture was incubated for 20 min on ice. Fixed bacteria were centrifuged for 30 min at 4°C at 2000 x g, and bacterial pellets were frozen at -80°C. Thawed pellets were resuspended in 10 ml of ChIP Buffer (1.1% Triton X-100, 1.2 mM EDTA, 16.7 mM Tris-HCl (pH 8.1), 167 mM NaCl, supplemented with cOmplete™Mini EDTA-free Protease Inhibitor Cocktail) and cells were lysed by 4 passages with French Press. The lysate was sonicated, and DNA size was checked with the 1.5% agarose gel. The lysate was centrifuged with the maximal speed and supernatant was collected. A 50 μl aliquot of supernatant was taken as “Input DNA” for the subsequent sequencing and data analysis. Following manufacturers’ protocols, remaining supernatant was immunoprecipitated ON with anti-FLAG Magnetic Agarose or anti-HA magnetic beads for 3xFLAG-tagged and HA-tagged proteins, respectively. Magnetic beads/agarose were immobilized and after removal of supernatant were sequentially rinsed with a panel of buffers: Low Salt Wash Buffer (0.1% SDS, 1% Triton X-100, 2 mM EDTA, 20 mM Tris-HCl (pH 8.1), 150 mM NaCl), High Salt Wash Buffer (0.1% SDS, 1% Triton X-100, 2 mM EDTA, 20 mM Tris-HCl (pH 8.1), 500 mM NaCl), LiCl Wash Buffer (250 mM LiCl, 1% NP-40, 1% deoxycholate, 1 mM EDTA, 10 mM Tris-HCl (pH 8.1)), TBS Buffer (50 mM Tris-HCl, 150 mM NaCl). The thawed Input control and immobilized Magnetic beads/agarose were mixed with 450 μl of TBS buffer, 15 µl of 5 M NaCl and 0.5 μl RNAse A (100mg/ml). The mixtures were incubated at 65°C ON. Then 4 µl of 0.5 M EDTA and 2 µl of 10 mg/ml Proteinase-K were added and the mixtures were incubated at 45°C for 2 h. The resulting mixtures were thoroughly mixed in 1:1 ratio with phenol/chlorophorm/isoamylalcohol combination and centrifuged at >10 000 x g for 1 min. The upper aqueous phase was transferred to a new tube, mixed in 1:1 ratio with chloroform and centrifuged at >10’000 x g for 1 min. The upper aqueous phase was transferred to a new tube and mixed with: (i) 25 μg of linear polyacrylamide; (ii) NaAc to a final concentration of 0.3 M; (iii) in 1:1 ratio with 100% isopropanol. The solution was centrifuged at >10’000 x g for 30 min, washed twice with cold 70% EtOH. Pelleted DNA was dried and resuspended in water ON. Library preparation and sequencing were done at the Genomics Facility Basel, D-BSSE, ETH-Zurich.

### Bioinformatics analysis

All alignments were done with the Clustal Omega and MUSCLE implemented in Geneious Prime 2020.2.3 (Edgar, 2004; Sievers et al., 2011). Mapping of ChIP-Seq data was done with bowtie2 (Langmead and Salzberg, 2012). Converting SAM to BAM, extracting unique binding reads and sorting were done with SAMtools (Danecek et al., 2021). Coverage plots were generated with bamCoverage and visualized with the Integrative Genomics Viewer (Ramírez et al., 2016; Robinson et al., 2011). Promoter predictions were done with BProm (Solovyev et al., 2011). Models of protein structures were either done with AlphaFold2 or taken from AlphaFold Protein Structure Database (Jumper et al., 2021; Varadi et al., 2022). Comparing the predicted models to Protein Data Bank (PDB) was done with the Dali server (Holm, 2022). Predictions of the secondary structure of the putative terminator was done with Mfold with the default parameters (Zuker, 2003). Calculations were performed at sciCORE (http://scicore.unibas.ch/) scientific computing centre at the University of Basel.

## Supplementary Figures

**Suppl. Figure 1.**
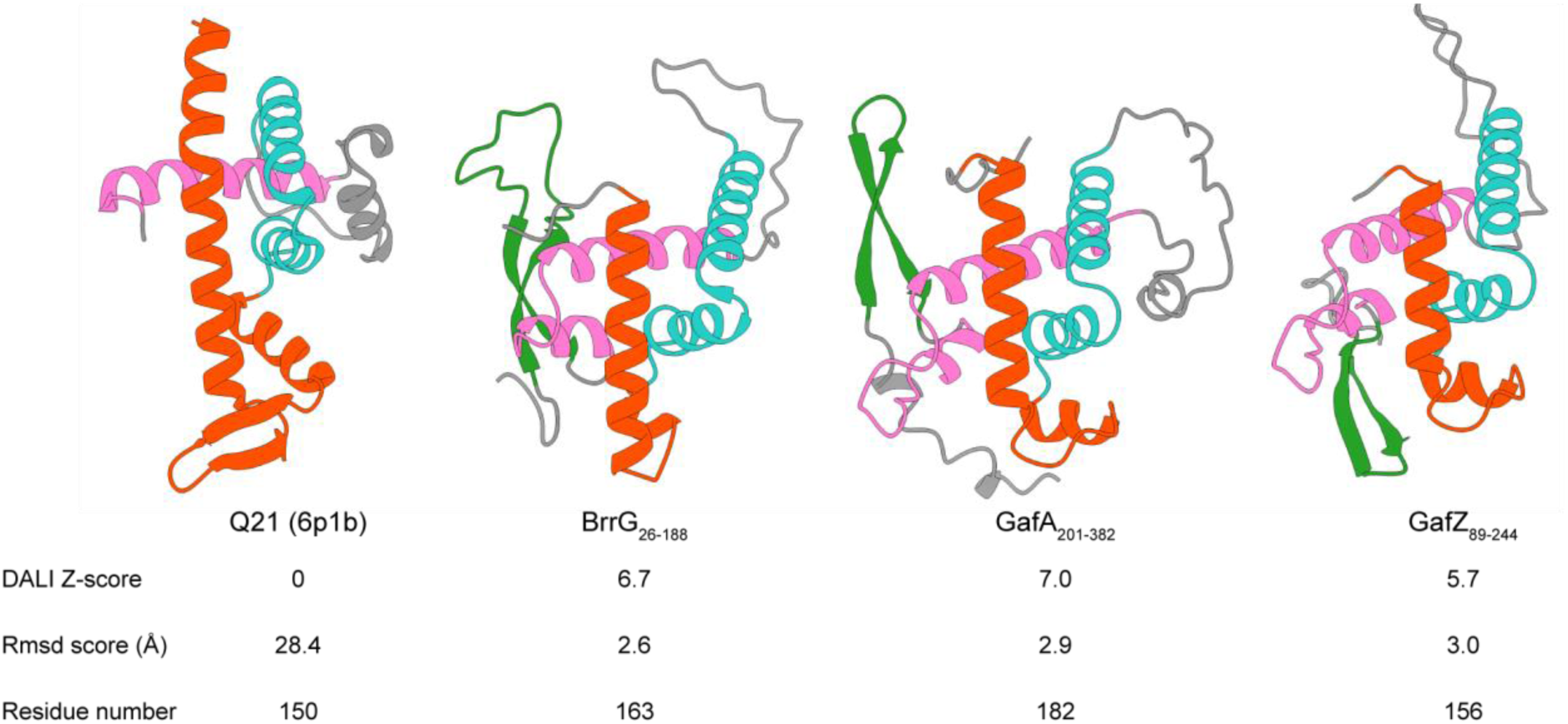
BrrG, GafA and GafZ share structural features. Structure of Q21 (6p1b) and AlphaFold2 models of BrrG (partial, A0A0R4J952), GafA (partial, D5AUH1) and GafZ (partial, A0A0H3C763). Similar features are shown in the same colour. Table below illustrates DALI Z-score and Rmsd score analysis of structural similarities. AlphaFold2 models were aligned to structure of Q21 (6p1b) using DALI Server (http://ekhidna2.biocenter.helsinki.fi/dali/). Higher Z-scores, respectively lower Rmsd values suggest better alignment and similarity to the reference structure.

**Suppl. Figure 2.**
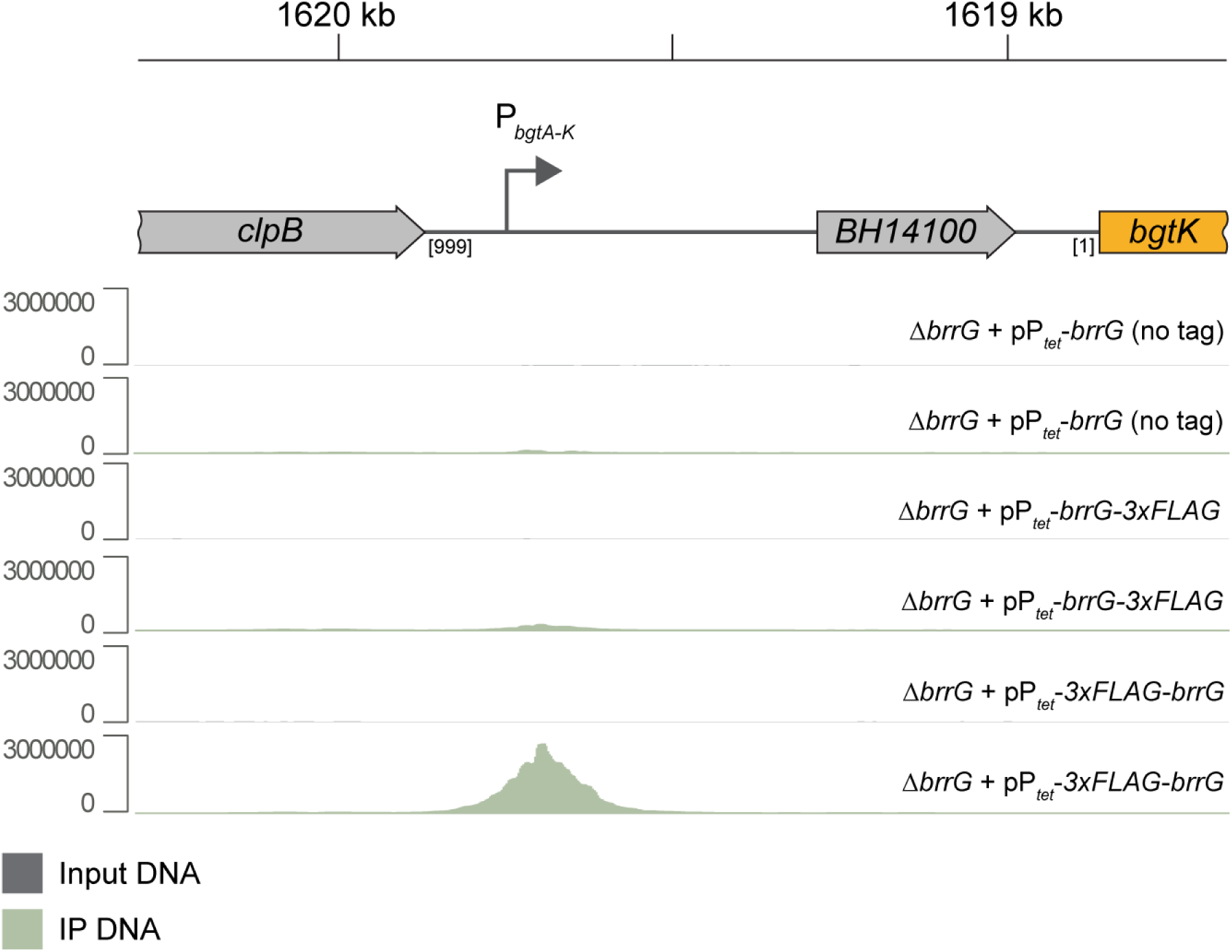
BrrG binds in the promoter region P*_bgtA-K_* in front of the *bgtA-K* cluster. Location of the P*_bgtA-K_* promoter and of the BrrG binding site of *ΔbrrG* + pP*_tet_-3xFLAG-brrG-* identified by ChIP-Seq. IP DNA is shown in green and input DNA in dark grey. *ΔbrrG* + pP*_tet_-brrG* was used as control. One replicate per strain is shown. Scale on the left indicates the coverage of the enriched DNA. Scale on the top indicates the genomic position in the *Bartonella henselae* genome.

**Suppl. Figure 3.**
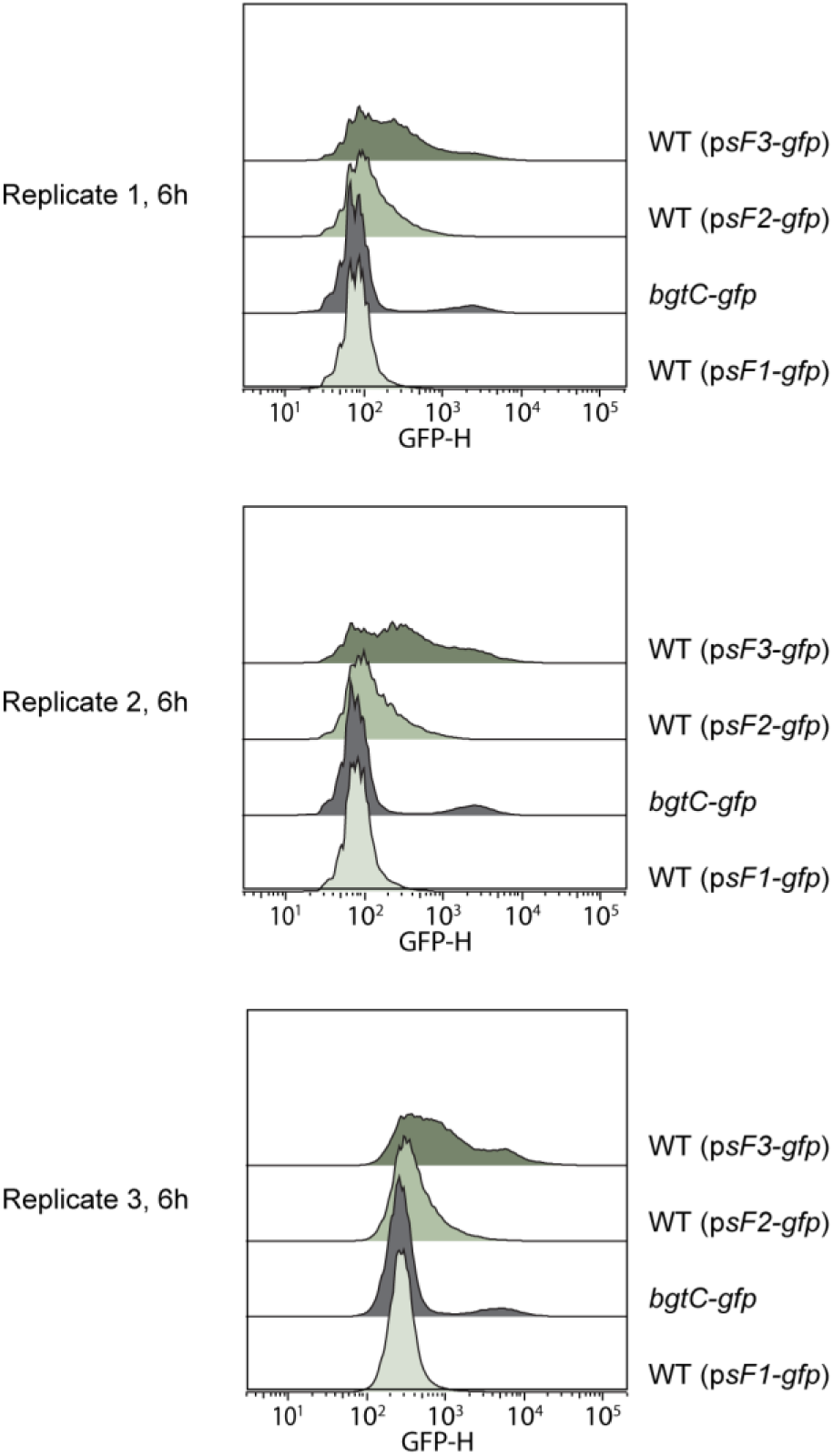
The P*_bgtA-K_*promoter is located far upstream of the *bgtA-K* locus. Fluorescence distribution at the 6 h time-point. Strain *bgtC-gfp* displaying wild-type expression pattern of the *bgtA-K* locus was used as control. The time-course experiments were started with the incubation of CBA agar-grown bacteria in M199 medium supplemented with 10% FCS. The experiments were done in biological triplicates.

**Suppl. Figure 4.**
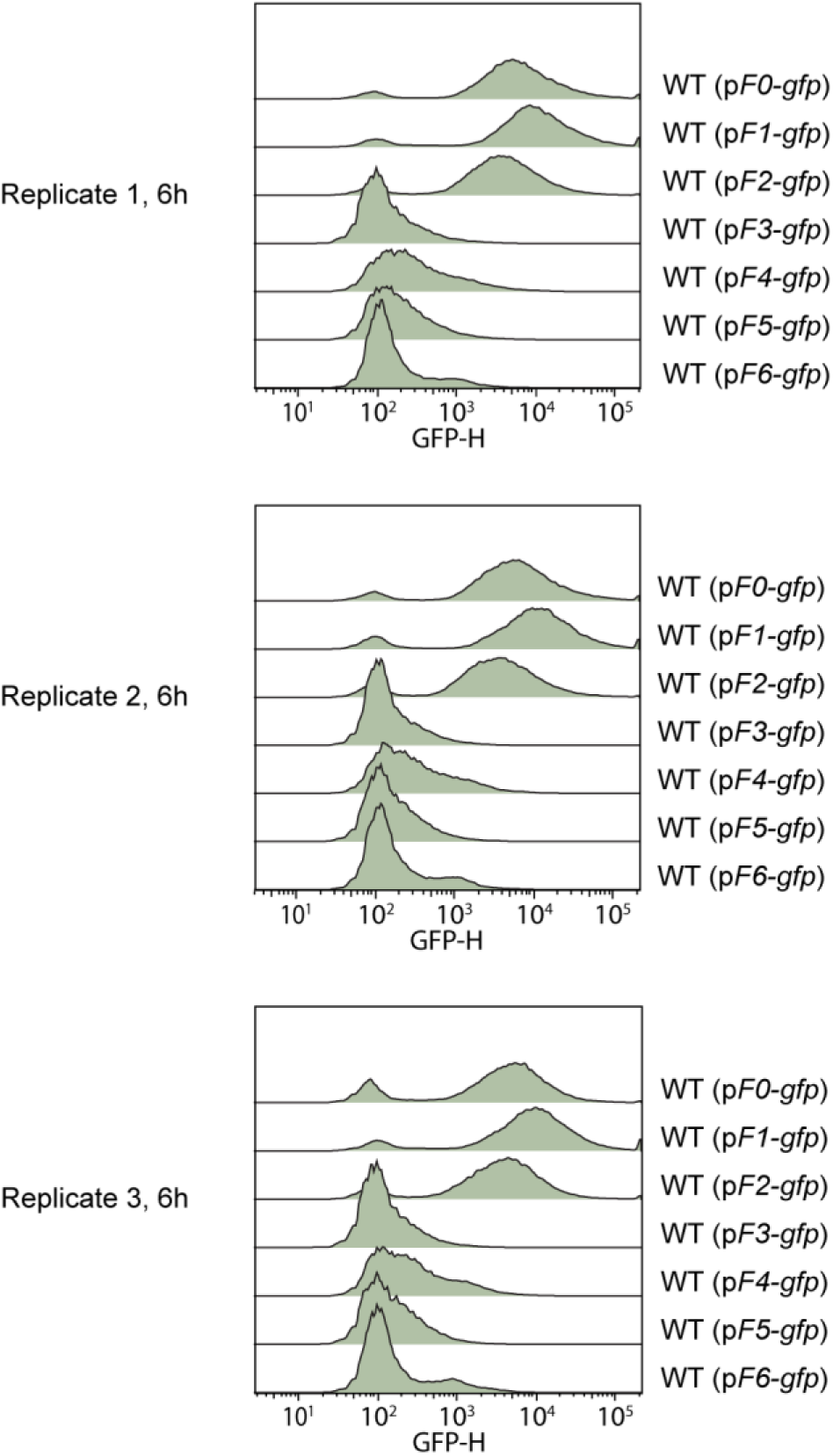
Fluorescence distribution for the bacterial populations containing the promoter-probe plasmids with various truncations at 6 h. The time-course experiments were started with the incubation of CBA agar-grown bacteria in M199 medium supplemented with 10% FCS. The experiments were done in biological triplicates.

**Suppl. Figure 5.**
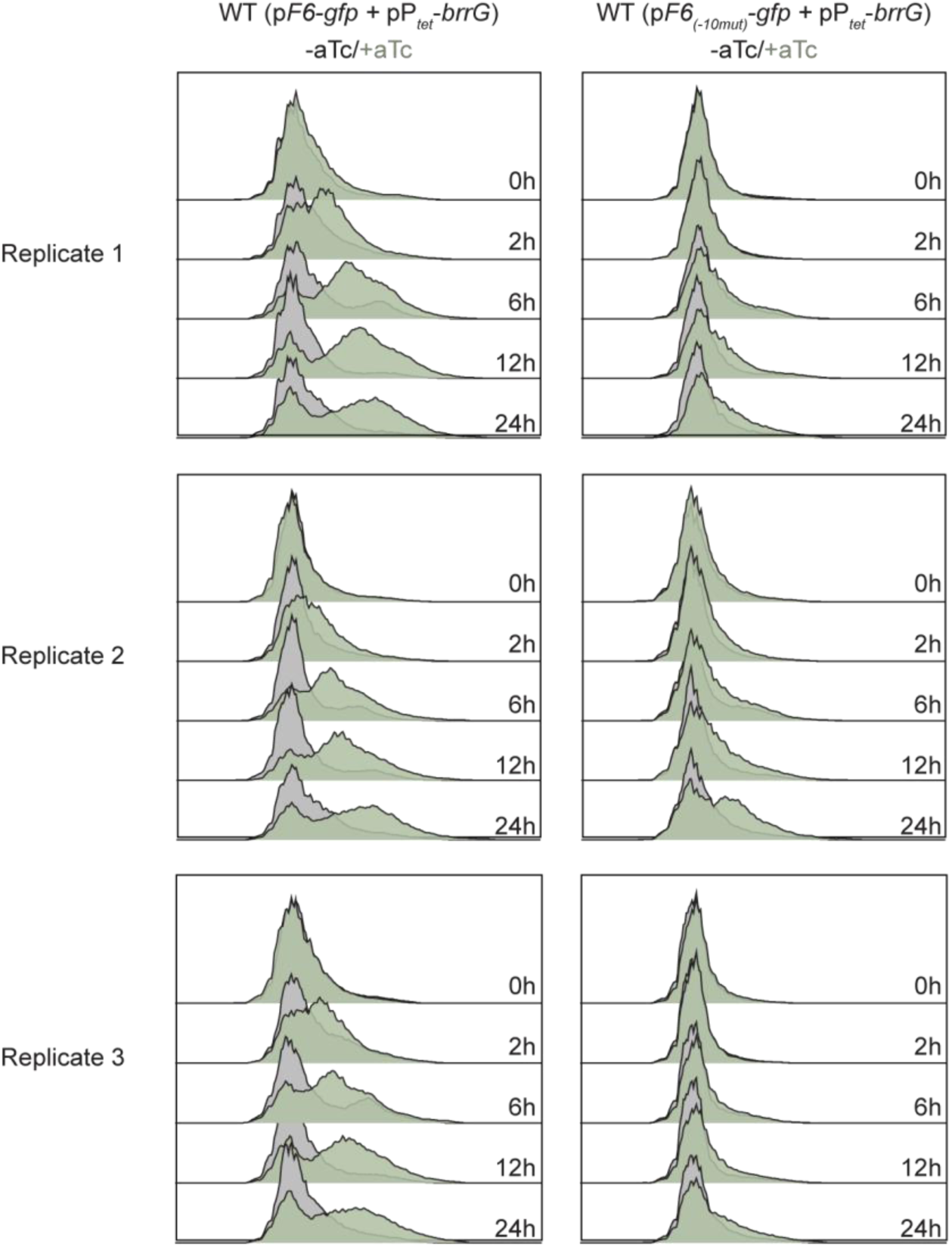
The -10-like element downstream of the P*_bgtA-K_* promoter is crucial for the antitermination by BrrG. Histograms of fluorescence distribution for the bacterial populations containing the promoter-probe plasmid pF6 with the P*_bgtA-K_* promoter and the adjacent native - 10-like element or its derivative, plasmid p*F6_(-10mut)_*, carrying mutations in two conserved positions of the -10-like element. Plasmid pP*_tet-brrG_* encodes an aTc-inducible *brrG* expression cassette. BrrG overproduction was induced by 25 ng/ml aTc. The time-course experiment was started with the incubation of CBA plate-grown bacteria in M199 medium supplemented with 10% FCS. Inducer aTc was added at time-point 0 h. Experiments were performed in biological triplicates.

**Suppl. Figure 6.**
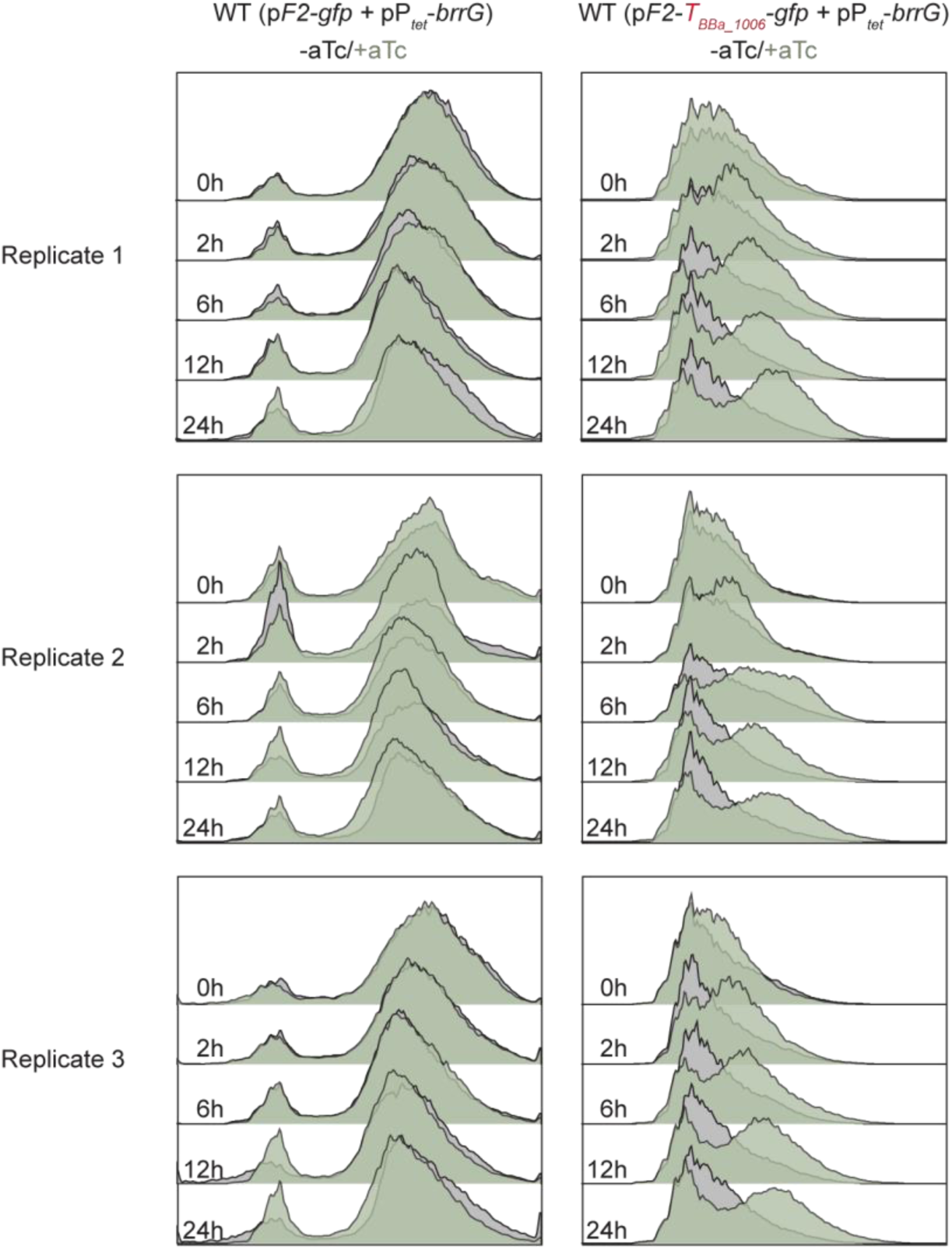
Fluorescence distribution for the bacterial populations containing the fragment F2 (Fig. 2A), the fragment F2 with the terminator and the sequence with the mutated -10-like element (pF6_-10mut_). Time of the measurements is indicated. Induction was done with 25 ng/ml aTc. Bacteria were cultivated in M199 medium supplemented with 10% FCS and aTc inducer at indicated concentrations. Experiments were performed in biological triplicates.

## Acknowledgement

We thank the FACS Core Facility of the Biozentrum for allowing us to conduct our FACS experiments and Dr. Jaroslaw Sedzicki for analysis for the structural analysis and figure preparation of the structure models. This work was supported by the Swiss National Science Foundation (SNSF, www.snf.ch) grant 310030B_201273 (to C.D.)

